# Connexin 43 Hemichannels Regulate Mitochondrial ATP Generation, Mobilization, and Mitochondrial Homeostasis against Oxidative Stress

**DOI:** 10.1101/2022.08.05.502934

**Authors:** Jingruo Zhang, Manuel A. Riquelme, Rui Hua, Francisca M. Acosta, Sumin Gu, Jean X. Jiang

**Affiliations:** Department of Biochemistry and Structural Biology, University of Texas Health Science Center, San Antonio, Texas, USA

**Keywords:** connexin 43, ATP synthase, proton, osteocytes, oxidative stress

## Abstract

Oxidative stress is a major risk factor that causes osteocyte cell death and bone loss. Prior studies primarily focus on the function of cell surface expressed Cx43 channels. Here, we reported a new role of mitochondrial Cx43 (mtCx43) and hemichannels (HCs) in modulating mitochondria homeostasis and function in bone osteocytes under oxidative stress. In osteocyte MLO-Y4 cells, the translocation of Cx43 to mitochondria was increased under H_2_O_2_-induced oxidative stress. H_2_O_2_ increased the mtCx43 level accompanied by elevated mitochondrial Cx43 HC activity determined by dye uptake assay. Cx43 knockdown (KD) by CRISPR-Cas9 lentivirus system resulted in impairment of mitochondrial function, primarily manifested as decreased ATP production. Cx43 KD had reduced intracellular reactive oxidative species (ROS) levels and mitochondrial membrane potential. Additionally, live-cell imaging results demonstrated that the proton flux was dependent upon mtCx43 HCs because its activity was specifically inhibited by an antibody targeting Cx43 C-terminus. The co-localization and interaction of mtCx43 and ATP synthase subunit F (ATP5J2) were confirmed by fluorescence resonance energy transfer (FRET) and a protein pull-down assay. Together, our study suggests that mtCx43 hemichannels regulate mitochondrial ATP generation by mediating K^+^, H^+^, and ATP transfer across the mitochondrial inner membrane and the interaction with mitochondrial ATP synthase, leading to enhancing the protection capacity of osteocytes against oxidative insults.

## Introduction

Connexins (Cxs) family consists of 20 members in mice and 21 in humans (Bedner et al., 2012; Söhl and Willecke, 2004). Six Cx subunits form a hexamer as a hemichannel (HC) with the ability to allow molecules less than 1 kDa to pass through (Goodenough and Paul, 2003; Jiang et al., 2007). HCs function as a gate for molecular communication between intracellular and extracellular spaces, allowing the passage of molecules and the exchange of information across the membrane. Two HCs in adjacent cells can form a gap junction, which assists in cell-cell communication directly from the cytoplasm of adjoining cells, thus synchronizing cellular functions (Iyyathurai et al., 2013). Under normal physiological conditions, undocked Cx HCs remain mainly closed (Sáez et al., 2010). However, pathophysiological conditions, such as injury or disease, drive their opening, leading to the augmentation of certain pathological conditions such as increased inflammatory reactions (Peng et al., 2022). HC opening is stimulated by many factors like low extracellular Ca^2+^ concentration, oxidative stress, mechanical signals, and extracellular alkalinization (Burra and Jiang, 2011; Decrock et al., 2011; Francis et al., 1999; Gault et al., 2014; Kar et al., 2012; Leithe et al., 2018; Schalper et al., 2010). The opening of Cx43 HCs also exhibits a positive impact on cells, especially in bone osteocytes (Kar et al., 2013; Zhao et al., 2022). Connexin 43 (Cx43) is the most ubiquitous expressed Cx in different cell types such as endothelial cells, cardiomyocytes, astrocytes, etc. It is also expressed in cellular organelles like mitochondria (Fiori et al., 2014; Gu et al., 2006; Li et al., 2002). Mitochondrial Cx43 (mtCx43) is localized with the C-terminus facing the mitochondrial intermembrane space (Miro-Casas et al., 2009). The physiological role of Cx43 in mitochondria remains largely elusive. Some data suggest that mtCx43 regulates K^+^ influx to the mitochondrial matrix, increasing mitochondrial respiration, ATP production, and reactive oxygen species (ROS) generation in cardiomyocytes (Boengler et al., 2012; Miro-Casas *et al*., 2009). The protective role mitochondrial Cx43 plays in pre-conditioning may be benefited from the channel property of Cx43 HCs. Many ion channels located in mitochondria are involved in ensuring the ion exchange between mitochondrial membranes. Ions, including Ca^2+^, Na^+^, K^+^, H^+^, Cl^-^, etc., not only participate in preserving normal mitochondrial membrane potential (ΔΨm), but are also involved in regulatory pathways (O’Rourke, 2007). Thus, the ion-permeable channels are very important to normal mitochondrial function, especially ATP production. In mitochondria, the permeability of mtCx43 HCs for ions and molecules is dedicated to a mild uncoupling of the mitochondrial inner membrane, which eventually causes protection for the ischemia/reperfusion (I/R) process (Ruiz-Meana et al., 2007).

Osteocytes are the most abundant cell in bone, occupying 90-95% of adult total bone cells (Bonewald, 2011). Cx43 HCs, richly present in osteocytes, prominently mediate anabolic action of mechanical loading and regulate the response to mechanical loading and oxidative stress (Cherian et al., 2003; Hua et al., 2021; Zhao *et al*., 2022). Oxidative stress causes apoptosis, further affecting the normal function of the osteocytic network. Cx43 HC is reported to reduce the cell death induced by oxidative stress in osteocytic MLO-Y4 cells (Kar *et al*., 2013). Plasma membrane Cx43 HCs open under oxidative stress, which protects osteocytes from apoptosis. Oxidative stress induces the accumulation of membrane Cx43 HCs at the cell surface, alleviating the oxidative stress caused by the oxidant stimulation. (Kar *et al*., 2013). The majority of prior studies focused on Cx43 HCs on the plasma membrane, while how mtCx43 function in osteocytes under oxidative stress remains largely unexplored. This study provides new mechanistic insight into the role of mtCx43 and HCs in regulating mitochondria function and protecting cells against oxidative damage.

## Results

### Cx43 translocates to mitochondria and forms functional HCs

To elucidate the role of Cx43 in osteocytes under oxidative stress, we treated osteocyte MLO-Y4 cells with H_2_O_2_ and investigated the cellular distribution of Cx43. We found that the mitochondrial localization of Cx43 was increased by its fluorescence signal overlap with succinate dehydrogenase (SDHA), a component of mitochondrial complex II (Figure 1A). Manders’ overlap coefficiency analysis indicated an increased co-localization between Cx43 and SDHA (Figure 1B). High magnification images further confirmed the overlap of SDHA and Cx43 in isolated mitochondria. SDHA and Cx43 fluorescence signals were merged in certain parts of mitochondria (yellow area) (Figure 1C). Western blot analysis further confirmed the significant accumulation of Cx43 protein in mitochondrial after 2 hrs of H_2_O_2_ treatment (Figure 1D) with a significant increase after 2 hrs of treatment, peaking after 3 hrs (Figure 1E). To assess if the mtCx43 formed functional HCs, a dye-uptake assay was performed. Lucifer yellow (LY) dye (425 Da), which is permeable by Cx43 HCs, was used as an indicator of active HCs, while TRITC-dextran (10 kDa), which is too large to pass through HCs, was used as a control to exclude non-specific dye uptake. The HC function was indicated by the ratio of LY/TRITC-Dextran. The dye uptake in isolated mitochondria, following H_2_O_2_, was elevated (Figure 1F). The treatment with carbenoxolone (CBX), a conventional Cx channel blocker, further confirmed that the increased dye uptake induced by H_2_O_2_ was caused by functional mtCx43 HCs. Since the C-terminus of mtCx43 faces the mitochondrial intermembrane space (Boengler et al., 2009; Miro-Casas *et al*., 2009), we also used the Cx43(CT) antibody, which specifically binds to Cx43 C-terminal to assess mtCx43 HCs. Cx43(E2) antibody, a specific Cx43 HC inhibitor targeting the extracellular loop of Cx43 as well as rabbit IgG were used as controls (Siller-Jackson et al., 2008). We found that in mitochondria isolated after H_2_O_2_ treatment, the dye uptake increment was inhibited by CBX and the Cx43(CT) antibody, while Cx43(E2) antibody had no effect on mtCx43 HCs (Figure 1F). The results suggest that Cx43 migrates to mitochondria, and mtCx43 HCs open in response to oxidative stress.

**Figure 1.**
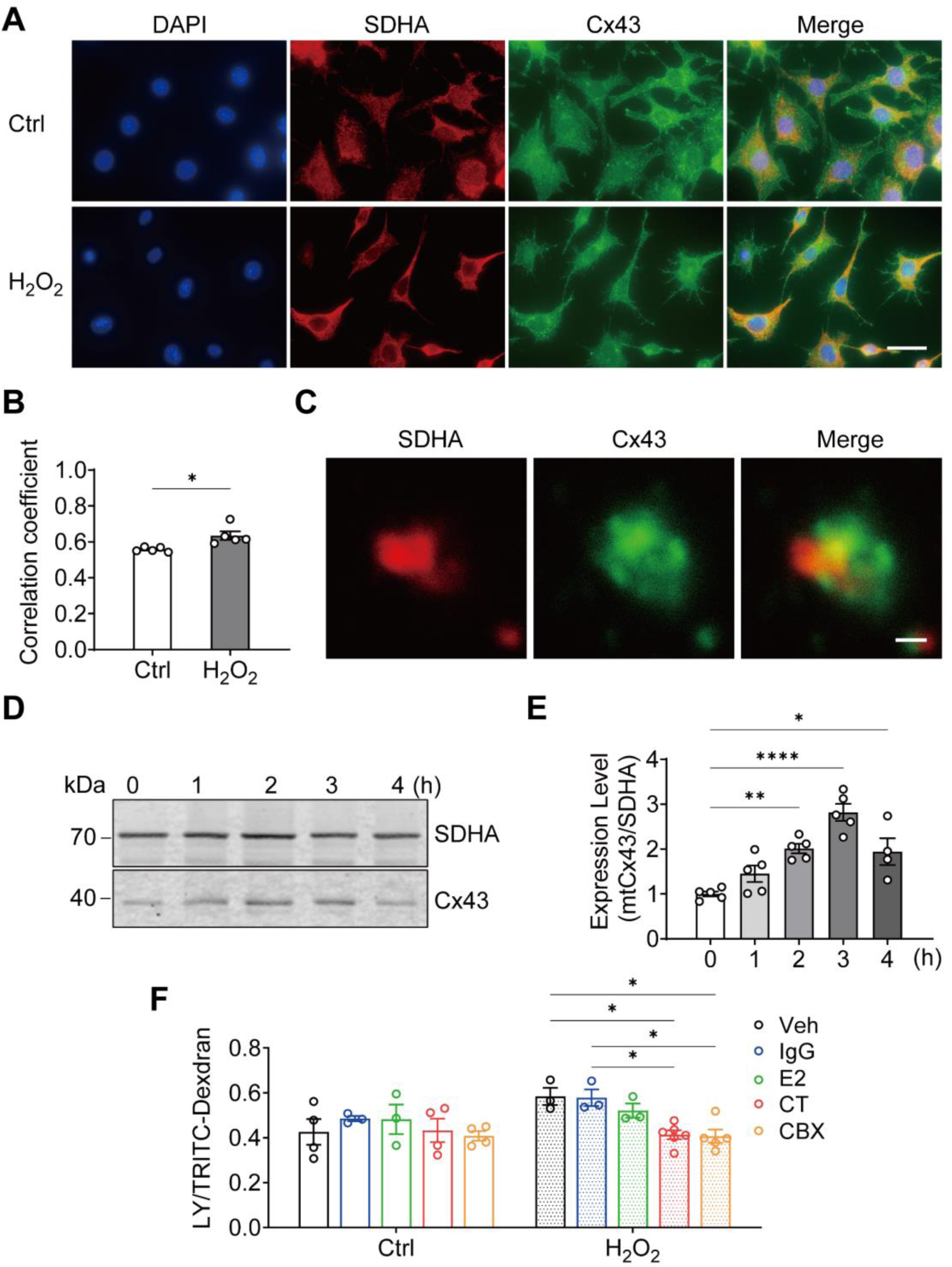
Cx43 translocated to mitochondrial and formed functional HCs induced by oxidative stress in osteocytes. **(A)** H_2_O_2_ treatment induced the translocation of Cx43 to mitochondria in the MLO-Y4 cells. MLO-Y4 cells were treated with 0.3 mM H_2_O_2_ for 2 hrs and fixed cells were immunostained with Cx43(E2) antibody and SDHA. Scale bar: 20 μm. **(B)** Manders’ overlap coefficient co-localization analysis of SDHA and Cx43 based on fluorescence signals. Data collected from 5 times independent experiments. Two-tailed t-test, \**p*<0.05. **(C)** Co-localization of SDHA and Cx43 in isolated mitochondria. Mitochondria were isolated from MLO-Y4 cells, fixed and subsequently immunostained with anti-Cx43 CT antibody and anti-SDHA antibody. Scale bar: 1 μm. **(D)** Western blot and statistical data showed a significant increase of Cx43 in isolated mitochondria after 0.3 mM H_2_O_2_ treatment. Isolated mitochondria from 0.3mM H_2_O_2_ treated MLO-Y4 cells during time were used for western blot analysis. \*\**p*<0.01. Data collected from N≥4 individual experiments. **(E)** Mitochondria dye uptake increased after H_2_O_2_ and inhibited by Cx43 C-terminal (CT) antibody and carbenoxolone (CBX). MLO-Y4 cells were treated with 0.3 mM H_2_O_2_ for 2 hrs for mitochondria isolation. Mitochondrial dye uptake assay was performed with LY/RD in the absence and presence of 0.5 μg/mL Cx43CT, Cx43(E2) antibody or 1μM CBX. N ≥ 3, Two-way ANOVA analysis was conducted, \**p*<0.05. TRITC-dexdran was used for calibration.

### Cx43 deficiency decreases mitochondrial membrane potential and ROS generation

By using the CRISPR-Cas9 lentivirus system, we knocked down (KD) Cx43 in MLO-Y4 cells with Rosa26 KD as a control. Whole-cell membrane and mitochondrial membrane proteins were extracted to evaluate the knockdown efficiency. An approximate 75% reduction in Cx43 protein level was seen from the total cell membrane (Figure 2A), and mitochondrial membrane protein (Figure 2B) extracts compared to the control group. Immunostaining of the isolated mitochondria from control and Cx43KD MLO-Y4 cells showed an overlap of SDHA and Cx43 proteins (orange signals), and the mtCx43 signal was much less in the Cx43KD group (Figure 2C). Meanwhile, the significantly increased dye uptake induced by H_2_O_2_ in control mitochondria was not shown in that of Cx43KD mitochondria (Figure 2D).

**Figure 2.**
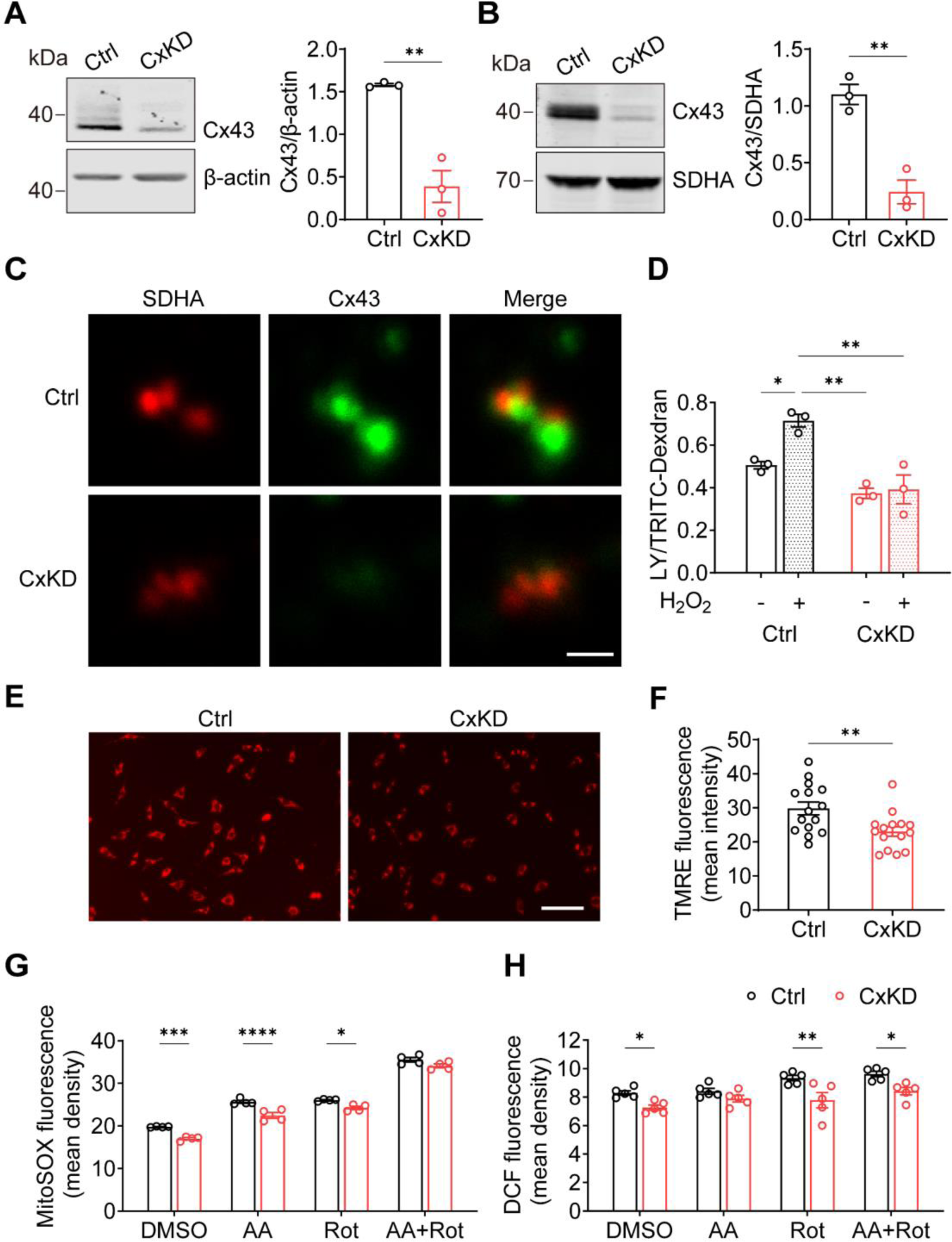
Cx43 knockdown reduced HC function in mitochondria. **(A)** Cx43 was significantly knocked down in MLO-Y4 cells by CRISPR-Cas9 system. Control and Cx43 KD (CxKD) MLO-Y4 cells were collected, and crude membrane extracts were analyzed using anti-Cx43 antibody. Band intensity was quantified and compared by Two-tailed t- test (right panel). N=3; \*\**p*<0.01. **(B)** Cx43 was significantly knocked down in isolated mitochondria. Mitochondria were isolated from control (Ctrl) and Cx43 knockdown (CxKD) MLO-Y4 cells and mitochondrial lysates immunoblotted with anti-Cx43 antibody. The KD efficiency in isolated mitochondria was quantified by the band intensity and compared by Two-tailed t-test. N=3; \**p*<0.05, \*\**p*<0.01. **(C)** Cx43 signal was absent in mitochondria of Cx43KD MLO-Y4 cells. MLO-Y4 cells with or without Cx43 knockdown were immunolabel with anti-Cx43 antibody and SDHA. Scale bar: 1μm. **(D)** Dye uptake in isolated mitochondrial Cx43 KD MLO-Y4 cells was abolished after H_2_O_2_ treatment. Mitochondrial were isolated from MLO-Y4 cells with 2 hrs 0.3 mM H_2_O_2_ incubation and LY/RD dye uptake assay was conducted and quantified. **(E)** Mitochondrial membrane potential (Δψm) was significantly decreased in the Cx43 KD MLO-Y4 cells. MLO-Y4 cells with or without Cx43 KD were incubated with TMRE which is used as an indicator for ΔΨm. **(F)** TMRE fluorescence signals were detected and quantified (right panel). N=5 of independent experiments. Two-tailed t-test was used for statistical analysis. Each experiment has triplicates with 3 images from each repeat. ** *p*<0.01. Scale bar:100 μm. **(G)** Cx43 KD significantly attenuated ROS production. MLO-Y4 cells with or without Cx43 KD were stained with mitoSOX and MitoSox fluorescence signals were quantified by NIH Image J and compared using Two-way ANOVA, N=4, * *p*<0.05, ** *p*<0.001. The cells were treated in the presence of complex inhibitors for 1hr. AA: Antimycin A, Rot: rotenone. **(G&H)** Mitochondrial ROS stained with DCF decreased in Cx43 impaired mitochondria. Mitochondria were isolated from either MLO-Y4 control or Cx43 KD cells. Treated with the inhibitors for 20 min. The fluorescence signals were measures using microplate reader and quantified by NIH Image J. Two-way ANOVA analysis and multiple comparitions was conducted in each group, N=5. * *p*< 0.05, ** *p*< 0.001. AA: Antimycin A, Rot: rotenone.

Mitochondrial electric membrane potential (ΔΨm), the main driver of oxidative phosphorylation (OXPHOS), is a key indicator of normal mitochondrial function.

Tetramethyl rhodamine and ethyl ester (TMRE) was used to measure ΔΨm, and its depolarization indicated a damaged mitochondrial inner membrane stabilization (Wang and Youle, 2009). The TMRE intensity was significantly reduced in Cx43KD cells, indicating the depolarization and impairment of ΔΨm (Figure 2E-F). To mitigate the possible effects of the cytosol, we used mitochondrial complex specific inhibitors antimycin A (AA) and rotenone (Rot) to stimulate the generation of mitochondrial ROS. We found that the basal level of mitochondrial ROS was already decreased after Cx43 KD compared with its parallel control. AA and Rot were capable of inducing ROS generation in both control and Cx43KD mitochondria. However, the Cx43KD group generated less mitochondrial superoxide with the existence of inhibitors both in intact cells, detected by mitoSOX (Figure 2G), and less ROS in isolated mitochondria, detected by DCF staining (Figure 2H). Together, the data suggest that Cx43 KD osteocytes had attenuated mitochondrial inner membrane potential and lower ROS generation.

### Mitochondrial coupling is impaired after Cx43 KD

Mitochondrial electron transport chain (ETC) coupling with oxidative phosphorylation is a fundamental mechanism for ATP generation by utilizing oxygen (Nolfi-Donegan et al., 2020). The decreased ΔΨm is likely to impede mitochondrial oxidative phosphorylation. We performed an ETC coupling study. With the application of plasma membrane permeabilizer (PMP) in the assay medium, we conducted measurements in sequential order: basal respiration (state 2), after the injections of ADP (state 3), oligomycin (OLIGO) (state 4o), FCCP uncoupler (state 3u), and antimycin A (AA) (Figure 3A-B). Oxygen consumption rate (OCR) in state 3 induced by ADP was significantly lower in Cx43 KD cells compared with the control group. The maximal respiration rate by FCCP uncoupling in state 3u was also lower in Cx43 KD cells compared to control. The non-mitochondrial OCR measured after injection of antimycin A (AA), a complex III inhibitor, was very low, almost to the same level as the control group. Both basal ATP production and ADP-induced ATP production were reduced in Cx43 KD cells. Proton leak, measured by subtracting OCR after antimycin A injection from state 4o, also showed a significant decrease in the Cx43KD group (Figure 3C). To demonstrate the ATP production level more directly, we extracted mitochondria and detected ATP amount in the entire cell, mitochondria, and cytosol. The bioluminescence assay based on ATP recombinant luciferase and the substrate luciferin were used for ATP detection. ATP concentration was much less in the whole-cell, mitochondria, and cytosol of Cx43KD compared with control cells (Figure 3D). The consistent data generated from mitochondrial coupling and ATP detection assays indicated the ATP production in Cx43 KD MLO-Y4 cells was remarkably decreased. In addition, quantitative PCR was conducted to further confirm if complexes in the mitochondrial respiration chain were affected at the mRNA level. We chose the marked proteins SDHA for complex II, ATP synthase subunit f (coded by *Atp5j2*) for complex V, and ADP/ATP translocase 2 (coded by *Slc25a5*) as its regulation of mitochondrial permeability transition pore (mPTP). The result showed the mRNA level of *Sdha* and *Atp5j2* was decreased in Cx43 KD cells compared to the control group, while that of *Slc5a5* did not show a significant difference (Figure 3E).

**Figure 3.**
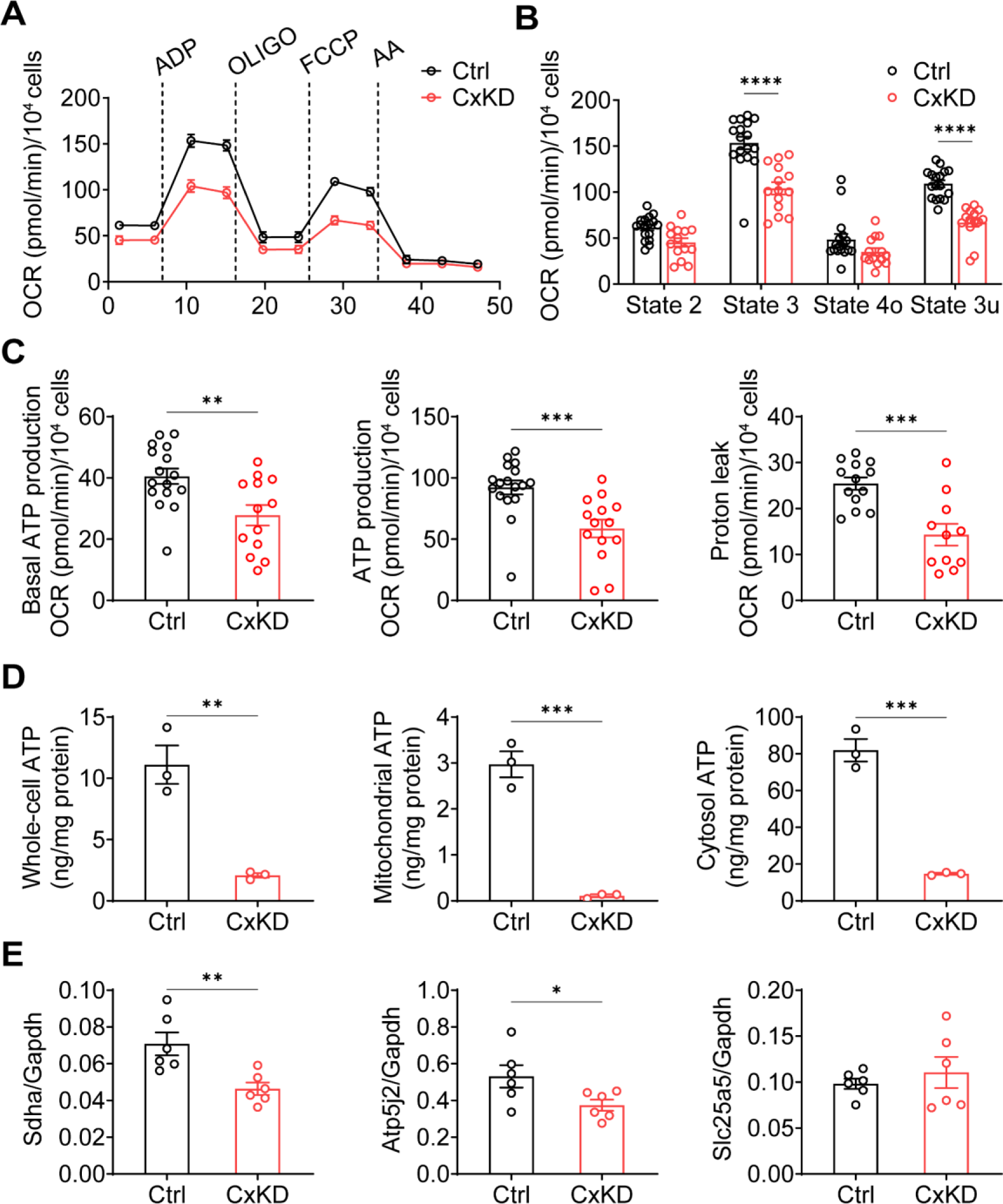
Mitochondrial functions were changed in the Cx43 KD Y4 cell line. **(A)** The mitochondrial oxygen consumption was impaired in Cx43 KD Y4 cells. Seahorse XF Cell Mito Stress Test was used to determine mitochondrial function in cells. Oxygen consumption rate (OCR) was measured in each state in which ADP, oligomycin (OLIGO), FCCP and antimycin A (AA) was used to treat cells. OCR level calibrated based on 10000 cells. **(B)** The OCR was significantly reduced after ADP and FCCP injections in Cx43 KD Y4 cells. \*\*\**p*<0.001. **(C)** The proton leak and ADP-induced ATP production, calculated by OCR, was decreased in Cx43KD experimental groups. N=3. N≥3 repeats in each experiment, \*\**p* <0.01, \*\*\**p*<0.001 **(D)** ATP was dramatically decreased in Cx43 KD Y4 cells. ATP determination was performed in isolated whole-cell lysis, isolated mitochondria, and cytosolic components. N=3. \*\**p* <0.01, \*\*\**p*<0.001 **(E)** The mRNA expression level of related genes in Y4 cell line. Mitochondrial complex II component protein succinate dehydrogenase (*Sdha*) and F(1)F(0) ATP synthase associated ATP synthase membrane subunit f (*Atp5j2*) were decreased in Cx43KD group. Mitochondrial permeable pore component ADP/ATP translocase 2 (*Slc25a5*) expression level had no significant changes between control and Cx43 KD Y4 cells. N=6, \**p* < 0.05, \*\**p*<0.01.

### Cx43 HCs increase the proton gradient across the mitochondrial inner membrane

mtCx43 is reported to modulate K^+^ influx into the mitochondrial matrix, and increased K^+^ in the matrix promotes the respiration and efflux of protons from matrix to intermembrane space (Boengler *et al*., 2012; Heinen et al., 2007). Proton gradient between intermembrane space and matrix is a key process for ATP generation. To determine the roles of mtCx43 HCs in proton gradient under oxidative stress, we used pHluorin, a GFP-based fluorescent protein that is sensitive to H^+^ signals, and live-cell imaging to detect proton changes. The process works by pH-insensitive blue fluorescent protein mTagBFP being fused before the N-terminus of pHluorin as a calibration, and the signal ratio of GFP/BFP increases during the increment of pH (Wu et al., 2019). mTagBFP-pHluorin was not specifically localized, primarily in the cytosol (Figure 4A). After H_2_O_2_ stimulation, a decrease in pHluorin signal was observed, while the signal change was significantly less in Cx43KD cells (Figure 4B). ATP5J2, acting as a subunit of ATP synthase, is localized on the mitochondrial inner membrane. We observed most of the ATP5J2-pHluorin signal was merged with mitoTracker Red (Figure 4-figure supplement 1). Therefore, the ATP5J2-pHluorin signal indicates the pH changes in the mitochondrial area. After inserting the *ATP5j2* sequence before the NH2-terminal of pHluorin, we monitored the pH signal (GFP/BFP) from ATP5J2-mTagBFP-pHluorin transfected MLO-Y4 cells (Figure 4C). H_2_O_2_ stimulation evoked a rise in the pHluorin signal in transfected cells (Figure 4D). This phenomenon was probably caused by the influx of protons from the mitochondrial intermembrane space to the mitochondrial matrix. The amplitude of the elevation was significantly lower in Cx43KD cells (Figure 4D). When plotting signal changes around the mitochondrial area, an oscillation of the mitochondrial pHluorin signal was observed. The amplitude of the oscillation was abolished in Cx43 KD mitochondria. Representative plots and the statistical analysis of the maximal amplitude of ATP5J2-pHluorin oscillation were shown (Figure 4E). Live-cell imaging results demonstrated that mtCx43 is critical for mitochondrial H^+^ flux in response to H_2_O_2_ stimuli.

**Figure 4.**
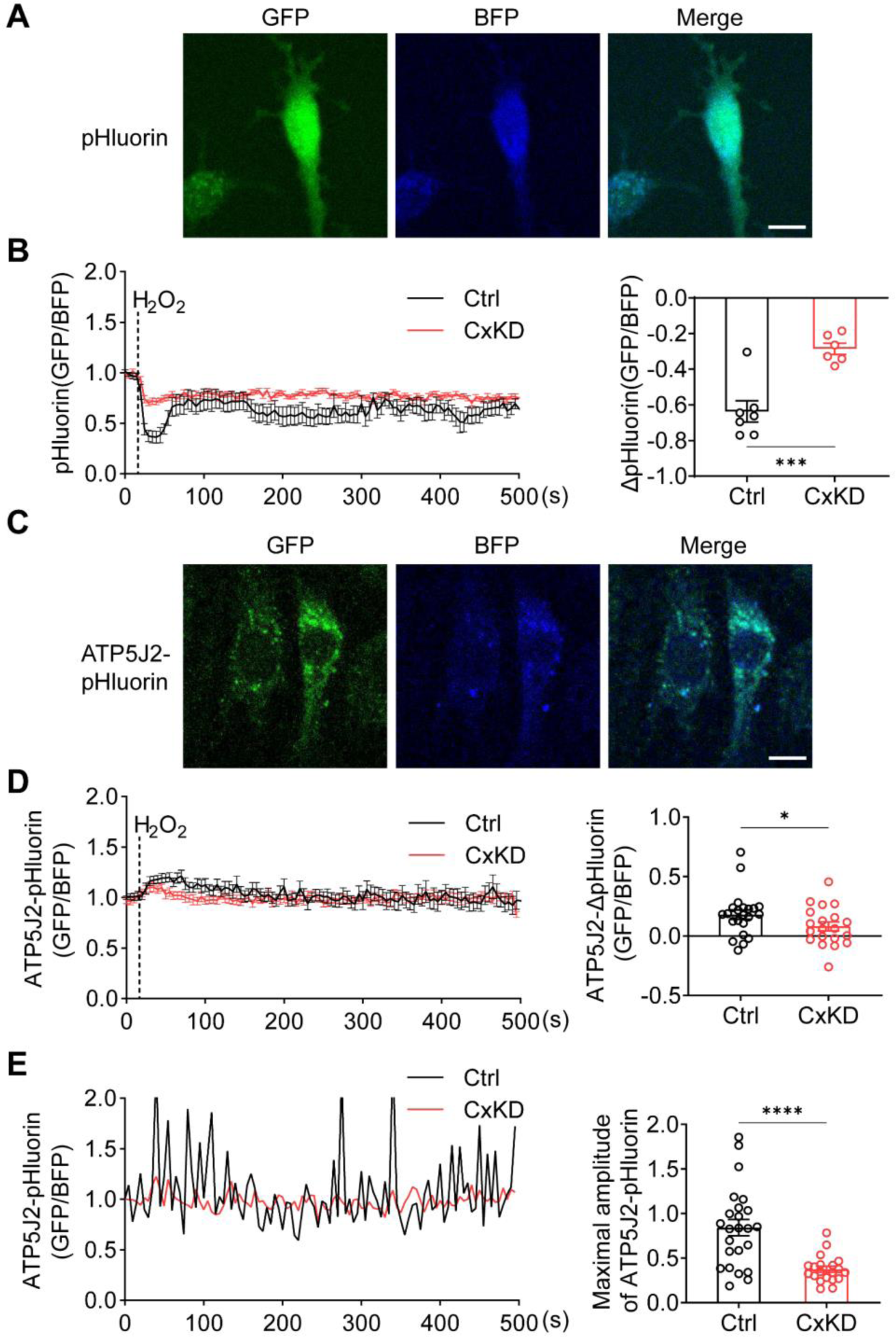
pHluorin signal indicated an attenuated sensitivity to oxidative stress in Cx43 KD Y4 cells. **(A)** pHluorin fluorescent protein localized in the cytoplasmic area in transfected MLO-Y4 cells. BFP is used as calibrations. **(B)** Cytosolic pHluorin signal change due to H_2_O_2_ stimuli was attenuated in Cx43 KD cells. Cytosolic pH change detected in Cx43 KD and control Y4 cells by pHluorin after 0.6 mM H_2_O_2_ stimuli at 25s time point. Statistical analysis showed the difference in the 40s time point. \*\*\**p*<0.001. **(C)** ATP5J2 conjugated pHluorin fluorescent protein localized on mitochondria. Green: pHluorin signal (GFP channel); Blue: BFP. Scale bar: 10μm. **(D)** Mitochondrial pHluorin signal change due to H_2_O_2_ stimuli was attenuated in Cx43 KD cells. ATP5J2-pHluorin signal indicated the mitochondrial pH level from Cx43 KD and control Y4 cells after 0.6mM stimuli at 25s time point. Statistical analysis showed the difference in the 40s time point in ATP5J2- pHluorin transfected Y4 cells. Data was collected from n≥7 cells in 3 times independent experiments. \**p*<0.05. **(E)** Mitochondrial pHluorin signal oscillation was inhibited in Cx43 KD Y4 cells. Representative ATP5J2-pHluorin signal oscillation in control and Cx43 KD Y4 cells. Two-tailed t test analysis showed significantly decreased oscillation amplitude in the CxKD group. \*\*\*\**p*<0.0001.

### Inhibition of mtCx43 HCs impedes mitochondrial function

Of importance is the membrane topology of Cx43 on the mitochondrial membrane, with its C-terminus facing intermembrane space, as we showed Cx43(CT) antibody, not Cx43(E2) antibody, has access to inhibit mtCx43 HCs (Fig. 1F). For this reason, we used the Cx43(CT) antibody as a specific mtCx43 HC inhibitor in the mitochondrial seahorse coupling assay after the addition of a plasma membrane permeable reagent (PMP). Cx43(CT) antibody treatment inhibited the increase of ATP production, similar to CBX, while the rabbit IgG control group did not show a reduction in oxygen consumption (Figure 5A). Respiratory control ratios (RCR, state 3/state 4o) and uncoupling control ratios (UCR, state 3u/state 4o) were similar among the groups without any significant difference (Figure 5B). The basal respiration (state 2) had no significant difference detected. State 3 after ADP stimulation and state 3u after FCCP uncoupling demonstrated lower OCR in the presence of CT antibody and CBX (Figure 5C). We further examined complex I activity using pyruvate and malate as substrates. CT antibody and CBX treated group showed the inhibitory role compared with Vehicle (Veh) and IgG control groups (Figure 5D). CT antibody, and CBX showed lower OCR, indicating reduced complex II activity (Figure 5E). The inhibitory role that CT and CBX played was mainly observed in Cx43KD MLO-Y4 cells in state 3. ADP-stimulated ATP production significantly decreased in CBX treated group; the CT antibody-treated group displayed a trend of inhibition while no significance was detected (Figure 5-figure supplement 1).

**Figure 5.**
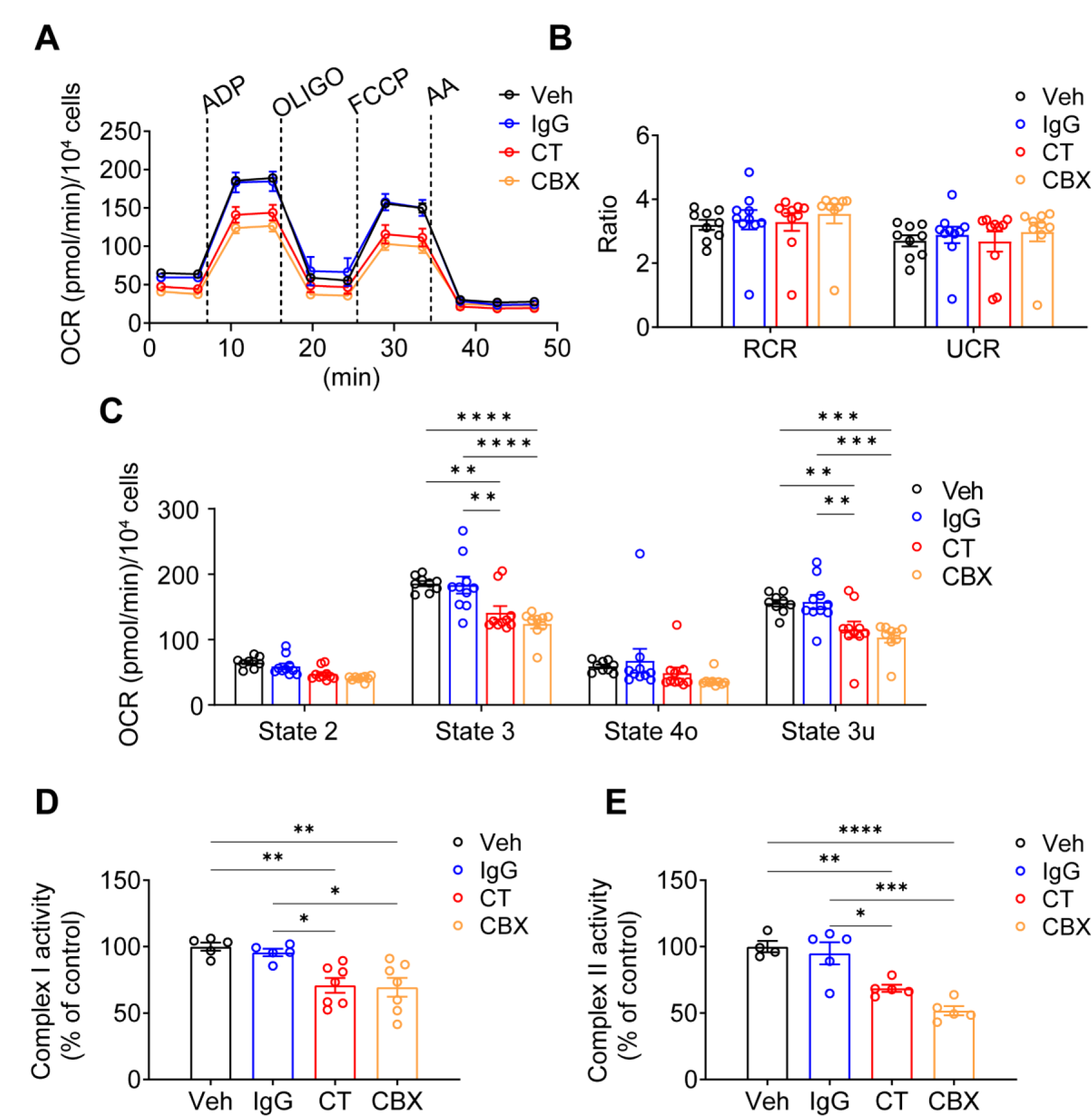
Cx43CT Ab treatment decreased mitochondrial respiratory capacity in MLO-Y4 cells. **(A)** Oxygen consumption rate (OCR) was measured on permeabilized MLO-Y4 cells, using an XF96e Analyzer. The sequential injection of mitochondrial inhibitors was indicated by arrows. OLOGO, oligomycin; AA, antimycin A. Rabbit IgG, Cx43CT antibody, CBX were pre-added in the assay medium. Basal respiration (state 2), after the injections of ADP (state 3), oligomycin (state 4o), FCCP (state 3u). **(B)** The respiratory control ratio RCR (RCR=state 3/state 4o) and UCR (UCR=State 3u/state 4o), which reflects the mitochondrial respiratory capacity, had no difference between groups. one-way ANOVA test. **(C)** OCR was detected significantly difference in state 3 and state 3u. Values corresponding to the different respiratory states are represented as mean ± SEM (n=15–18 replicate of three independent experiments/group). Two-way ANOVA analysis was performed. \**p*<0. 05, \*\*\**p*<0. 001, \*\*\*\**p*<0. 001. **(D)** Both CT antibody and CBX treated group showed the inhibitory role on complex I activity. Complex I activity was measured in Xfe96 Seahorse bioanalyzed by sequential administration of a combination of pyruvate (10 mM)/malate (0.5 mM), FCCP (0.5μM), and antimycin A (4μM). The activity was normalized to the control group. \**p*<0. 05, \*\**p*<0. 01. **(E)**. CT antibody and CBX treated group showed the similar inhibitory role on complex II activity. Complex II enzymatic activity was analyzed similarly to Complex I. After basal OCR measurements cells were sequentially treated with a combination of rotenone (2μM)/succinate (10 mM), FCCP (0.5μM), and antimycin A (4μM). All data were reported as the mean ±S.E. \**p*<0. 05, \*\**p*<0. 01, \*\*\**p*<0. 001, \*\*\*\**p*<0. 001.

To further validate the involvement of mtCx43 in proton changes, we generated a Cx43-pHluorin plasmid and expressed it in MLO-Y4 cells (Figure 6A). The Cx43- pHluorin signal in live imaging was quantified in the whole cell as well as the local mitochondrial area after H_2_O_2_ treatment. A decrease in whole-cell Cx43-pHluorin signals calibrated with BFP was observed after adding H_2_O_2_ (Figure 6B). In the mitochondrial area, indicated by mitochondrial marker mitoTracker Red, a similar oscillation of pHluorin signal was demonstrated (Figure 6C). Since Cx43 was expressed in different subcellular regions, the whole-cell and mitochondrial signals showed consistent pHluorin signals in the cytosol and ATP5J2-pHluorin signals in mitochondria. PMP was used to permeate the plasma membrane with intact mitochondria. With a permeable plasma membrane, the Cx43(CT) antibody could reach and function in mitochondria. Our results showed the CT antibody caused a reduction of pHluorin signal oscillation when compared with IgG treatment (Figure 6D-E). Although there was a minor effect, the CT antibody did not show significant inhibition of ATP5J2-pHluorin signals (Figure 6-figure supplement 1). The results that the inhibition of mtCx43 HCs by Cx43(CT) antibody caused a decreased pHluorin signal indicated the reduced proton gradient across the mitochondrial inner membrane. The increased pH gradient by mtCx43 HCs may be one of the underlying mechanisms that impacts mitochondrial function in response to oxidative stress. Considering that the proton gradient across the mitochondrial inner membrane is essential for ATP synthesis, we investigated the connection between mtCx43 and mitochondrial ATP synthase.

**Figure 6.**
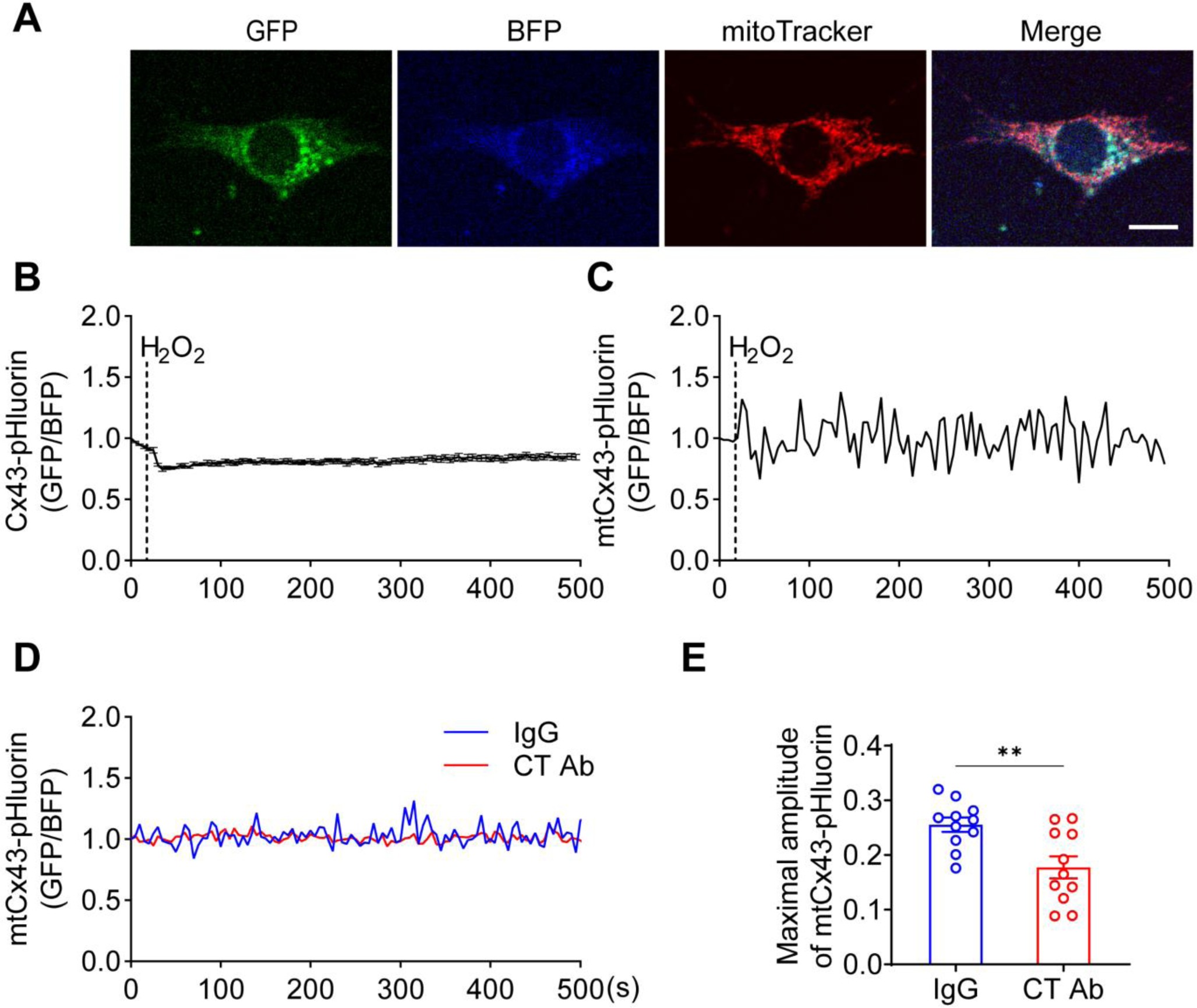
MtCx43 HCs were permeable to protons indicated by Cx43-pHluorin signal. **(A)** Typical images of Cx43-pHluorin in MLO-Y4 cells. Cx43-pHluorin fluorescent protein localization in transfected MLO-Y4 cells. Green: pHluorin signal (GFP channel); Blue: BFP; Red: mitoTracker Red. Scale bar: 10μm. **(B)** Cx43-pHluorin signal was reduced in whole-cell area after 0.6mM H_2_O_2_ treatment. BFP signal was used for calibrations**. (C)** Representative mitochondrial Cx43-pHluorin oscillation in MLO-Y4 cells after 0.6mM H_2_O_2_ treatment. BFP signal was used for calibrations. **(D-E)** CT antibody reduced Cx43- pHluorin signal oscillation. Mitochondrial pHluorin signal oscillation was recorded in Cx43-pHluorin transfected MLO-Y4 cells. The maximal amplitude of the oscillation was analyzed. \*\**p*<0. 01.

### Cx43 directly interacts with ATP synthase subunit ATP5J2

We showed that mtCx43 HCs are functionally involved in ATP generation. We then determined the spatial relationship between mtCx43 and ATP generation machinery mitochondrial complex V. In complex V, ATP5J2 is the subunit that circles the ATP synthase peripheral stalk with the NH2-terminus facing the matrix and a membrane inserted helix (He et al., 2018), and was identified to be associated with Cx43 in human chondrocytes (Gago-Fuentes et al., 2015). To investigate if there is physical interaction between the mtCx43 and ATP synthesis protein, we employed the Förster resonance energy transfer (FRET) approach. Cx43-cyan fluorescent protein (CFP) and ATP5J2-yellow fluorescent protein (YFP) were co-transfected into MLO-Y4 cells. CFP and YFP form a FRET pair in which CFP works as the donor and YFP as the acceptor (Miyawaki et al., 1997). The FRET signal was detected when the two fluorescent proteins were in proximity of less than 10 nm to each other. ATP5J2-YFP signal was primarily detected in the mitochondria region, while the Cx43-CFP signal was also detected in other parts of the cell. However, the FRET signal was detected (Figure 7A), suggesting that Cx43 and ATP5J2 physically interacted with each other in the mitochondria. An immunoprecipitation assay was also conducted using the GFP or ATP5J2-GFP transfected MLO-Y4 cells (Figure 7-figure supplement 1A). Cx43 band was detected when an anti-GFP antibody against ATP5J2-GFP was used as an immunoprecipitating antibody (Figure 7B). To further investigate the binding between Cx43 and ATP5J2 in mitochondria, we performed a protein pull-down assay with ATP5J2-GFP transfected MLO-Y4 cell extracts using GST- Cx43 CT immobilized on glutathione-agarose beads (Figure 7-figure supplement 1B). The results showed that GST-Cx43 CT, not GST control, was able to pull down ATP5J2-GFP, further demonstrating a direct interaction of Cx43 CT with ATP5J2 in intermembrane space (Figure 7C). Together, these findings suggested a direct binding of complex ATPase subunit ATP5J2 with Cx43 CT in mitochondria.

**Figure 7.**
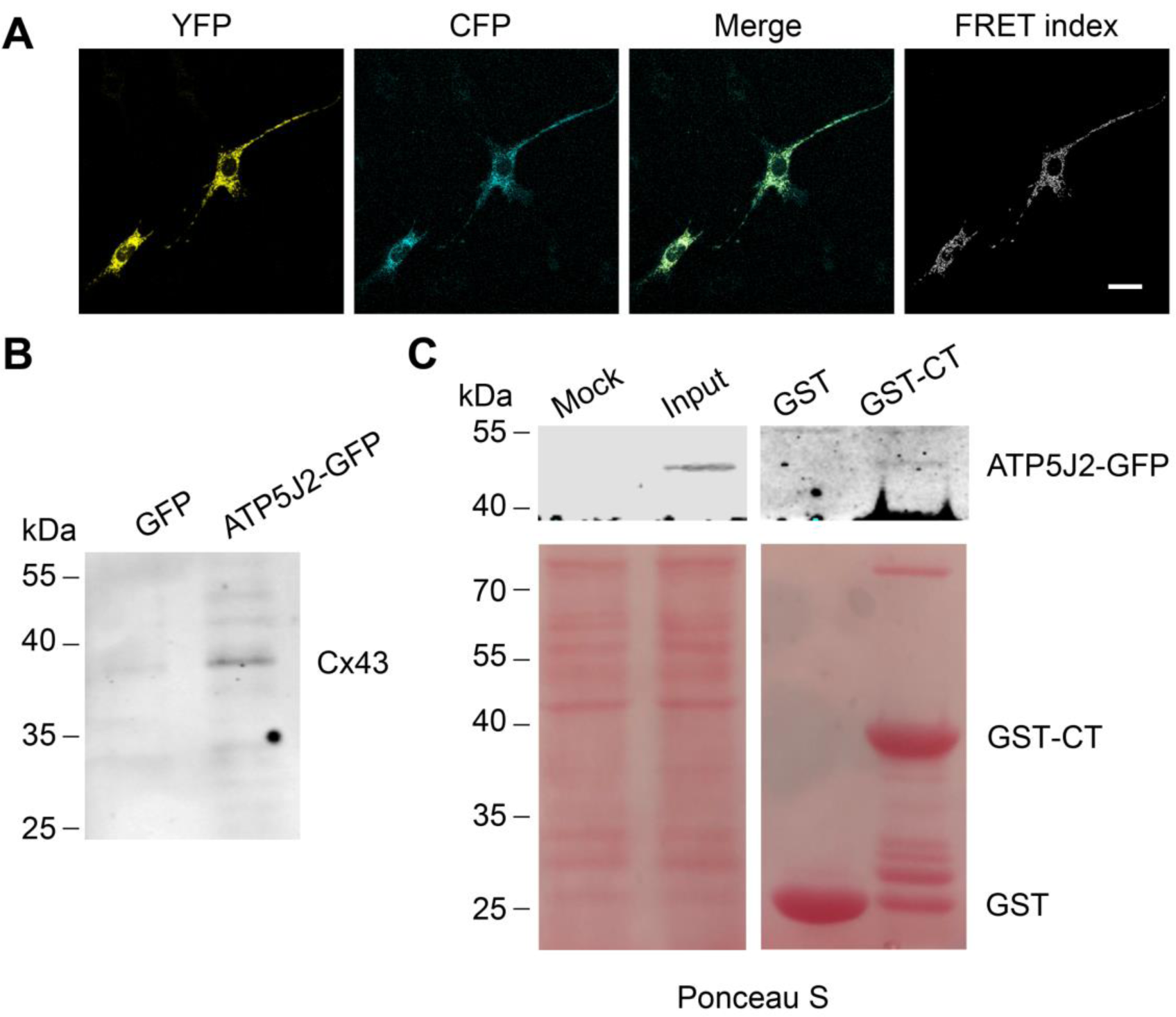
Cx43 interacted with ATP5J2 on mitochondria. **(A)** FRET assay showed the interaction between Cx43 and ATP5J2 in MLO-Y4 cells. MLO-Y4 cells co-expressing Cx43-CFP and ATP5J2-EYFP were presented. EYFP fluorescence (EYFP was excited, and its fluorescence recorded; the first column), CFP fluorescence (CFP was excited and its fluorescence recorded; the second column), and “FRET index” (images processed with the “FRET and Colocalization Analyzer”, fourth column). Scale bars: 20 μm. **(B)** Immunopaticification showed physically binding of Cx43 with endogenous ATP5J2. Cx43 band was detected after immunoprecipitation using ATP5J2-GFP as the bait. After IP, samples were collected and ran an SDS-PAGE. Anti-Cx43 (1:300) was used to incubate the membrane. **(C)** GST pull-down assay proved the physical binding between ATP5J2, and Cx43-CT. Purified GST-CT was conjugated to glutathione agarose beads with GST used as control.

### Cx43 accumulated in mitochondria of the oxidized osteocytes in *Csf-1^+/-^* mice in vivo

To assess whether Cx43 migrates and accumulates in mitochondria in response to oxidative stress *in vivo*, we used a transgenic mouse model with increased oxidative stress in osteocytes. Our previous study suggested that deficiency of macrophage-colony stimulating factor 1 (CSF-1), a key factor involved in cell signaling and remodeling, increases Nox4 oxidase expression in primary osteocytes, resulting in elevated oxidative stress (Werner et al., 2020). We isolated the primary osteocytes from *Csf-1^+/-^* mice and measured the ROS level by DCF staining. We observed a significantly increased ROS production in primary osteocytes with decreased CSF-1 (Figure 8A). The significant increase of co-localization of Cx43 and SDHA suggested the increased migration of Cx43 to the mitochondrial in *Csf-1^+/-^* osteocytes compared with the WT (*Csf-1^+^*^/*+*^) cells (Figure 8B). The result recapitulated the mitochondrial migration and accumulation of Cx43 in oxidized osteocytes *in vivo*.

**Figure 8.**
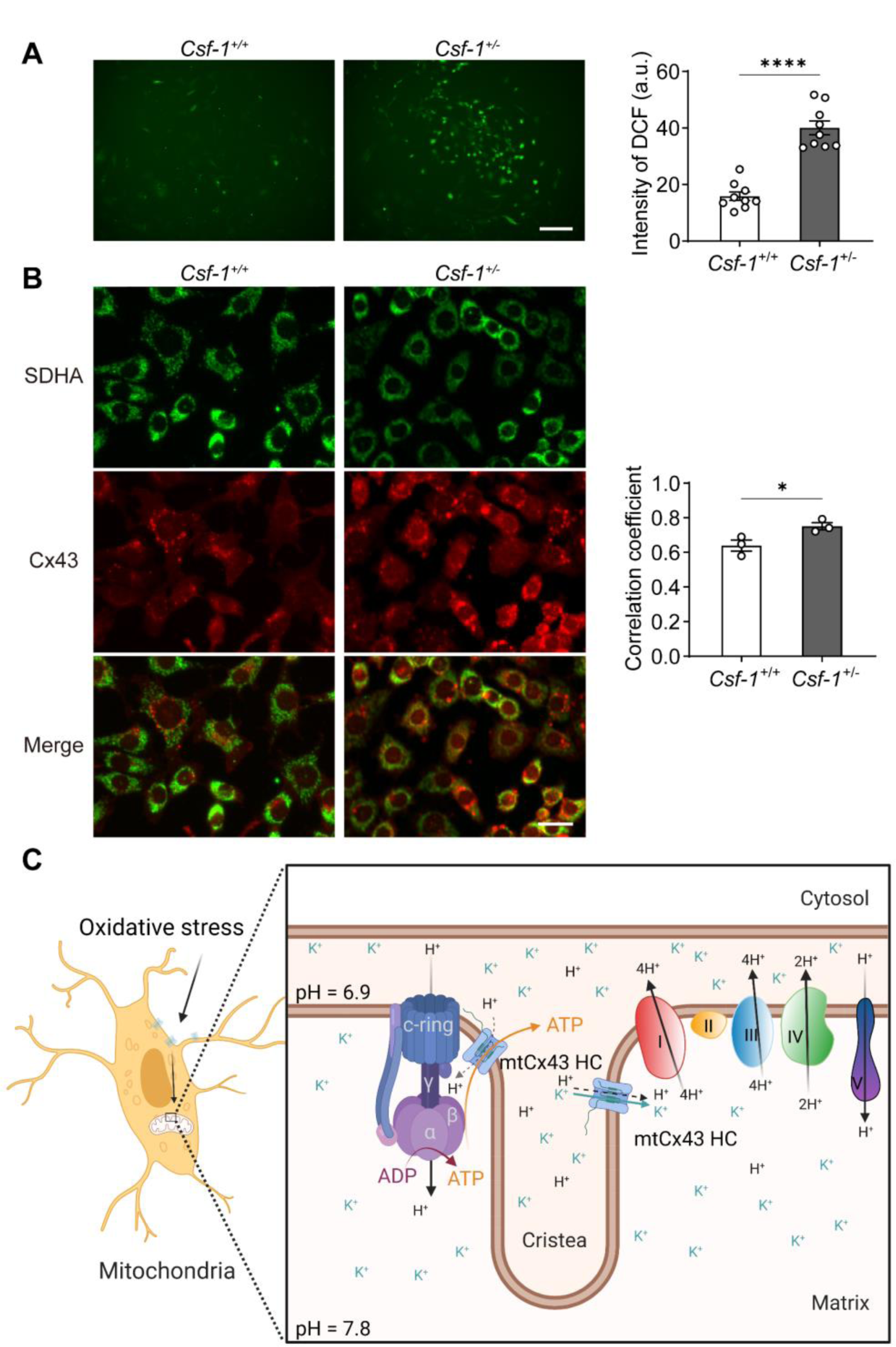
mtCx43 response to oxidative stress in primary osteocytes. **(A)** Osteocytes from *Csf-1^+/-^*mice have an increased ROS level by DCF staining. Data collected from 3 animals in each group, each point indicated the ROS level from each individual well. Scale bar: 50 µm. Two-tailed t-test was conducted, *\*\*\*\*p<0.0001.* **(B)** Osteocytes from *Csf-1 ^+/-^* mice have an increased co-localization of SDHA (Green) and Cx43 (Red). Primary osteocytes were stained with SDHA and Cx43 antibody. Scale bar: 20 µm. Coefficient co- localization analysis of SDHA and Cx43 based on fluorescence signals. Data collected from 3 mice each group. Two-tailed t-test, *p<0.05. **(C)** Schematic diagram on mtCx43 function in osteocytes. In oxidative stress, Cx43 is translocated onto mitochondria. With a special position close to ATP synthase, mtCx43 HCs enabled the H^+^ influx between mitochondrial inner membrane, accelerating ATP production. Thus, mtCx43 HCs play a crucial role in osteocytes under oxidation.

## Discussion

Our study showed that accumulated Cx43 in mitochondria played an important role in protecting mitochondria and cell homeostasis agonist oxidative stress. mtCx43 participates in mitochondrial membrane potential. mtCx43 HCs participate in proton gradient maintenance between intermembrane space and matrix, enhancing mitochondrial ATP generation. Moreover, the direct physical interaction between mtCx43 and mitochondrial ATP5J2 demonstrated the coupling of mtCx43 with a subunit of ATP synthase, the mitochondrial complex V. This interaction is likely to have a positive impact on transferring ATP generated by complex V via mtCx43 HCs to increase ATP supply in the cytosol, a critical step for protecting cells against oxidative stress. Together, our data unveiled a new mechanism concerning the unique role of mt Cx43 HCs in mitochondria function, ATP generation, and transport under oxidative insults.

Osteocytes, as the most abundant bone cell types, are dynamic and responsive cells, forming a network in the skeleton to orchestrate bone modeling and remodeling (Bonewald, 2011). Oxidative stress is a major insult to bone tissue, and increased oxidative stress during postmenopausal or glucocorticoid medication is one of the major underlying causes of bone loss and osteoporosis (Domazetovic et al., 2017). Here, we observed that when an acute oxidant stimulus occurs, Cx43 is rapidly translocated into the mitochondria inner membrane. We used Cx43 KD MLO-Y4 cells to explore the role of mtCx43 and found that mtCx43 expression was tightly correlated with ATP generation. Decreased oxygen consumption rate in permeabilized Cx43 KO MLO-Y4 cells indicated the role of Cx43 in the coupling of the mitochondrial inner membrane and ATP production. Mitochondria are the fuel factory for cells, which produce bioenergetic source ATP for cellular activities (Spinelli and Haigis, 2018). Mitochondrial electron transport complexes I-V are responsible for ATP production (Zhao et al., 2019). Accompanied by transporting electrons, mitochondrial electronic complexes generate a proton gradient between mitochondrial intermembrane space and mitochondrial inner matrix (Perry et al., 2011). ATP synthase, also known as complex V, utilizes this proton gradient to produce ATP from ADP, eventually preserving the normal homeostasis and function of cells.

Given the vital role of protons in mitochondria, we utilized the pHluorin as the proton indicator to investigate the impact of mtCx43 on local proton activity indicated by pH changes. Cx43 KD MLO-Y4 cells had reduced pH change in both cellular and mitochondrial compartments. It has been reported that plasma Cx43 channels permeate protons (Swietach et al., 2007). If mtCx43 HCs mediate proton transfer between intermembrane space and matrix, we would expect to observe the increased proton gradient in Cx43 KD cells. Instead, we observed a decreased ATP5J2-pHluorin signal and pH gradient in Cx43 KD cells. The reduced proton level in the intermembrane space and pH gradient was also detected when mtCx43 HCs were inhibited by Cx43(CT) antibody. Cx43 has four transmembrane domains with a long cytosolic CT. Due to mtCx43 orientation with CT facing mitochondrial intermembrane space (Boengler et al., 2009; Miro-Casas et al., 2009), Cx43(CT) specific antibody, other than Cx43(E2) antibody, functioned as the gating inhibitor for mtCx43 HCs. The addition of PMP that permeabilizes plasma membrane permits the entry of the Cx43(CT) antibody into the cell and targets the mtCx43 in intact mitochondria. The consistent results obtained with Cx43 KD and treatment with Cx43(CT) antibody also reduced protons at the intermembrane space and the pH gradient. These results indicate that mtCx43 HCs may not primarily act as a leaky proton channel, but instead may regulate the ion transport that indirectly increases the proton transport against the proton gradient. Indeed, the mitochondrial membrane potential measured by TMRE was decreased in Cx43 KD cells.

In mitochondria, both H^+^ and ^·^OH gradients across intermembrane space are important for the mitochondrial membrane potential (Lee, 2019). If taking the small pH chemical gradient into account, the mitochondrial membrane potential is, more correctly, a protonmotive force (Nicholls, 2002). The electric membrane potential of mitochondria ranges from -150 ∼180 mV (Santo-Domingo and Demaurex, 2012). Wide-range changes of H^+^ influx will cause the depolarization of the mitochondrial inner membrane. The opening of mitochondrial permeability transition pores (mPTP) causes a propagated loss of mitochondrial membrane potential. Meanwhile, mPTP opening can be triggered by Ca^2+^ and ROS (Ong and Gustafsson, 2012). mtCx43 is reported to engage in the inhibition of mPTP opening, ultimately resulting in the maintenance of ATP production (Ferko et al., 2019). Azarashvili et al. also revealed that CBX shortened the lag time before mPTP opening and accelerated the Ca^2+^-induced permeability transition in synaptic and nonsynaptic rat brain mitochondria (Azarashvili et al., 2010). In adipose tissue, specific knockdown of Cx43 leads to a significantly lower expression of components on mitochondrial complex II, III, and V (Kim et al., 2017). The decreased associated proteins in electron transport chains may contribute to the lower energy expenditure.

Cx43 HCs have a large conductance (∼220 pS) with relatively low ion selectivity, allowing the passage of ions such as Ca^2+^, Na^+^, and K^+^ (De Smet et al., 2021; Wang et al., 2013). Previous studies indicated the permeability of mtCx43 HCs to K^+^ (Boengler et al., 2013; Miro-Casas *et al*., 2009). The submaximal K^+^ entry into the mitochondria matrix enhances electron flow with a preserved membrane potential and production of ROS (Heinen *et al*., 2007). To counter the K^+^ influx, mitochondrial electron transport chain complexes I, III, and IV pump protons out, into intermembrane space to enhance the proton gradient (Nolfi-Donegan *et al*., 2020). mtCx43 HCs, thus, are likely to mediate the transfer of K^+^ from the intermembrane space into the matrix. However, we cannot exclude the possibility of passage of protons following the gradient from the intermembrane space to the matrix. In intermembrane space, K^+^ concentration is about 140 mM, while H^+^ is around 0.1 μM (Feher, 2012; Garlid and Paucek, 2003). Therefore, the amount of K^+^ passed through mtCx43 HCs from intermembrane space to matrix driven by electrical gradient is expected to be proportionally much greater than that of H^+^. We did observe the reduction of proton leak in Cx43 KD mitochondria, indicting the relatively minor role of mtCx43 HCs in transferring H^+^ from intermembrane space. pHluorin is a pH-sensitive GFP variant, and the use of fusion protein by linking pHluorin with Cx43 CT permits the detection of proton changes close to Cx43 in the intermembrane space. The local Cx43-pHluorin signal in the mitochondria region displayed an oscillation pattern, suggesting that there is proton flux activity close to the mtCx43. Cx43(CT) antibody abolished this oscillation, suggesting mtCx43 HCs regulate proton flux in the intermembrane space. Taken together, the mtCx43 HCs are likely to play a predominant role in transferring K^+^ into matrix, thus the proton output into intermembrane space through electron transport complexes is more significant than proton influx into matrix; thus, increasing membrane potential, proton gradient, and ultimately generating more ATP.

The majority of proton leak across the mitochondrial inner membrane is thought to be correlated to the adenine nucleotide translocase (ANT) and uncoupling proteins (UCPs) (Divakaruni and Brand, 2011; Jastroch et al., 2010). Proton leak is tightly linked to ROS generation; increased mtROS generated from ETC induce proton leak, and proton leak suppresses ROS production as a feedback loop (Cheng et al., 2017). There are conflicting studies on proton leak and ROS generation; few reports suggest an increased ROS level after proton leak (Boveris and Chance, 1973; Starkov and Fiskum, 2003), while most studies prove the positive feedback loop of proton leak and ROS generation (Brookes, 2005; Echtay et al., 2002). The controversy may be caused by the amplitude of proton leak, and mild-moderate proton leak is reported to be protective during I/R injury in cardiomyocytes (Nadtochiy et al., 2006). In our case, the proton leak mediated by mtCx43 HCs appear to be at mild-moderated levels and thus, may play a protective role on osteocytes.

Under normal conditions, very fewer mtCx43 are present in the mitochondria, while under oxidative stress, the accumulation of Cx43 in the mitochondria inner membrane and functional mtCx43 HCs appear to play significant roles in enhancing the ATP generation as well as export of ATP into the cytosol. We showed a significant reduction of ATP in both mitochondria and cytosol with the deletion of Cx43 in osteocytes. Cx43 HCs have been reported to mediate the transport of ATP (Kang et al., 2008; Patel et al., 2014). Mass spectrometry analysis in mouse hearts indicated the new interaction between Cx43 with apoptosis-inducing factor (AIF) and the beta-subunit of the electron-transfer protein (ETFB). AIF and ETFB are typical proteins regulating mitochondria oxidative phosphorylation (Denuc et al., 2016). We showed that mtCx43 co-localized and interacted with a subunit of ATPase, ATP5J2 in complex V, and the ATP level in the cytosol was reduced in Cx43 KD. It is likely that mtCx43 HCs may enhance the transport of the freshly generated ATP by complex V to the inner membrane space and cytosol. Cx43 KD had no effect on the expression of *Slc25a5* gene, an ATP/ADP translocase, suggesting that increased ATP level in the cytosol is not caused by the impact of mtCx43 on this transporting mechanism. Thereby, the close physical interaction may help accelerate the export of ATP synthesized in the mitochondrial matrix by complex V. The increased ATP level in cytosol will protect the cells against oxidative damage since oxidative stress depletes cytosolic ATP, leading to cell apoptosis. mtCx43’s protecting role on a cardiac pre-condition (Rodríguez-Sinovas et al., 2018) may also benefit from these mechanisms.

Together, the translocation of Cx43 to mitochondria in osteocytes after oxidative stress acts as a cell “rescue mission” by promoting ATP production through increased mitochondrial membrane potential and proton gradient through activation of mitochondrial complexes. The localization and interaction of complex V ATPase may aid in the transfer of ATP into the cytosol in protecting cells against oxidative stress (Figure 8C).

## Materials and Methods

### Cell culture and reagents

MLO-Y4 osteocyte cells established from primary osteocytes were cultured on type-Ⅰ rat tail collagen coated dishes in α-modified essential medium with 2.5% fetal bovine serum (FBS) and 2.5% bovine calf serum (BCS) with penicillin-streptomycin (PS) (Kato et al., 1997). The KD of Cx43 by CRISPR-Cas9 system in MLO- Y4 cells with Rosa26 KD as a control was previously described (Hua et al., 2022). Transfected MLO-Y4 cells were cultured without penicillin-streptomycin. Cx43(E2) antibody and Cx43(CT) antibody were both developed in the Jiang laboratory (Cheng et al., 2001; Siller-Jackson *et al*., 2008). Cx43(E2) antibody has been used as a Cx43 HC inhibitor as previously described (Kar *et al*., 2013; Sáez et al., 2013; Siller-Jackson *et al*., 2008). Mouse anti-SDHA antibody was purchased from Invitrogen (#459200, Invitrogen, Waltham, MA, USA).

### Immunofluorescence

Cells were seeded in cell culture plates or 12-mm glass coverslips for 2 days before fixation with 4% paraformaldehyde (PFA) for 10 min at room temperature (RT). After washing in PBS buffer for 3 times, cells were incubated with affinity-purified antibodies against Cx43(E2) (1:300 dilution) or SDHA (1:1000 dilution) in blocking buffer (0.25% Triton X-100, 2% gelatin, 2% donkey serum, and 1% bovine serum albumin) for 1 hr at RT, followed by incubation with fluorescein-conjugated secondary antibody for 1 hr at RT. After incubation with 4, 6-diamidino-2-phenylindole (DAPI) for 5 min, washed the cells with PBST for three times. After mounting, the slides were ready for imaging using a fluorescence microscope (Keyence BZ-X710, Tokyo, Japan).

### Protein extraction and immunoblot analysis

Membrane protein extracts from MLO-Y4 cells were prepared for western blot and GST pull-down assays. Cells were lysed and homogenized by a 20G needle with ice-cold lysis buffer (5 mM EDTA, 5 mM EGTA and 5 mM Tris, pH 8.0). Proteinase inhibitors (2 mM PMSF, 5 mM NEM, 1 mM Na_3_VO_4_ and 0.2 mM leupeptin) were added. The cell lyses was then centrifuged at 600 × g for 5 min, removing intact cells and nucleis. Subjected the supernatant to centrifuge (Beckman Coulter, Brea, CA, USA) at 125,000 × g for 30 min at 4°C. The pellets were then resuspended in lysis buffer with 1% SDS for western blot or in 0.5% Triton X-100 buffer for GST pull-down assay.

Protein concentration was quantified using the micro-BCA assay (Pierce, Rockford, IL, USA), and equal amounts of proteins were loaded on SDS-polyacrylamide gel electrophoresis (PAGE) and electroblotted onto a nitrocellulose membrane. Membranes were incubated first with primary antibodies, anti-Cx43(CT) (1:300 dilution), anti-SDHA (1:1000, Invitrogen), and anti-β actin antibodies (1:5000) overngiht at 4℃, followed with the incubation of secondary antibodies, goat anti-rabbit IgG conjugated IRDye® 800CW or goat anti-mouse IgG conjugated IRDye® 680RD (1:15,000 dilution), 1 hr at RT. The images were captured using a Licor Odyssey Infrared Imager (Lincoln, NE, USA), and protein band intensity was quantified using NIH Image J software.

### DNA construct preparation and transfection

The coding region of mouse Cx43 (*Gja1)* was amplified by PCR and DNA fragments were inserted into the EGFP-N1 or CFP-N1 vector to generate Cx43-EGFP or Cx43-CFP, respectively. *Atp5j2* cDNA was inserted into the EGFP-N1, EYFP-N1 vector to generate *Atp5j2*-EGFP or *Atp5j2*-EYFP, respectively. The mTagBFP-pHluorin plasmid was used to generate *Cx43*-mTagBFP-pHluorin and *Atp5j2*- mTagBFP-pHluorin constructs. For live-cell imaging, MLO-Y4 cells were transfected with *Cx43*- mTagBFP-pHluorin or *Atp5j2*-mTagBFP-pHluorin using the Neon transfection system with parameters set as 1200 mV with a duration of 20 ms, 1 pulse. The ratio of cells and transfected plasmids was 10^6^ cells with 5 µg plasmids. The expression of endogenous plasmids was determined by fluorescence imaging intensity.

### Mitochondria isolation

MLO-Y4 cells were collected, washed with ice-cold PBS twice, and centrifuged for at 1000 × g for 5 min at 4°C. Resuspended the pellets in 1 ml of mannitol buffer (220 mM mannitol, 70 mM sucrose, 20 mM HEPES, 1 mM EGTA, 0.1% BSA, adjusted to pH 7.2 with KOH) and then homogenized. The homogenate was centrifuged at 1000 × g for 10 min to remove intact cells and nucleis, the supernatant was then centrifuged at 9000 × g for 15 min to precipitate down mitochondria. The pellet containing mitochondria was washed once with mannitol buffer and was resuspended in mannitol buffer with 1% BSA for further usage.

### Mitochondrial ROS measurement

Intracellular ROS was determined using mitoSOX (#M36008, Invitrogen, Waltham, MA, USA) in intact MLO-cells cells or using fluorescence-based probes Carboxy-H_2_DCFDA (#C2938, Invitrogen, Waltham, MA, USA) in isolated mitochondria. Cells were rinsed with Hanks’ balanced salt solution (HBSS) for 3 times and incubated with 100 nM mitoSOX for 20 min at 37°C. After three washes with HBSS, fluorescence images were immediately captured using a Keyence BZ-X710 microscope. Isolated mitochondria were stained with 10 μM Carboxy-H2DCFDA for 10 min at 37℃, and the data were collected using a microplate reader (Bio-Tek Synergy HT, Winooski, VT).

### Mitochondrial dye uptake assay

Mitochondria dye uptake assay was performed based on published studies (Miro-Casas *et al*., 2009; Soetkamp et al., 2014). Isolated mitochondria from MLO-Y4 cells were resuspended in isosmotic succinate buffer (150 mM KCl, 7 mM NaCl, 2 mM KH_2_PO_4_, 1 mM MgCl_2_, 6 mM MOPS, 6 mM succinate, 0.25 mM ADP, and 0.5 μM rotenone with pH 7.2) at 0.4 mg/mL. Mitochondria were then aliquoted into 5 groups, with the addition of either vehicles or Cx43 HC blockers. After a 20 min incubation period at RT and 200 × g centrifuge for 3 min, 50 μM of the Cx43 hemichannel-permeable dye Lucifer Yellow CH dilithium salt (LY, #L453, Invitrogen, Waltham, MA, USA) and 25 μg/mL of the hemichannel impermeable dye Tetramethylrhodamine-dextran 10,000 MW (TRITC-dextran, #D1816, Invitrogen, Waltham, MA, USA) were added. Incubated samples for 25 min and centrifuge at 200 × g at RT. Mitochondria were subsequently washed for 2 times, and gently resuspended in 200 μL succinate buffer. Fluorescence of LY (λ_ex_ 430 nm/λ_em_ 535 nm) and TRITC- dextran (λ_ex_ 545 nm/λ_em_ 600 nm) was measured by a microplate reader (Bio-Tek Synergy HT, Winooski, VT).

### Mitochondrial function assay

Mitochondrial oxygen consumption rates (OCR) were measured using a Seahorse Xfe96 Analyzer (Seahorse Bioscience, North Billerica, MA), which was equilibrated at 37°C overnight. 5X10^3^ MLO-Y4 cells were seeded in a pre-coated XF96 plate and cultured overnight for adherence. On the same day of the experiment, 50 μL containing 1 x mitochondrial assay solution (220 mM mannitol, 70 mM sucrose, 10 mM KH_2_PO_4_, 2 mM HEPES, 5 mM MgCl_2_, 1 mM EGTA, 0.2% BSA, adjusted pH 7.2 with KOH) with 10 mM succinate and 2 μM rotenone as substrates for the coupling assay was added to each well of the XF96 plate and followed by addition of plasma membrane permeabilizer (PMP, Invitrogen). To assess various parameters of mitochondrial function, OCRs were measured after injecting ADP, oligomycin (a complex V inhibitor), p-trifluoromethoxy carbonyl cyanide phenylhydrazine (FCCP, a protonophore and mitochondrial uncoupler), and antimycin A (a complex III inhibitor). The final concentrations of drugs in the well were ADP, 5 mM; oligomycin, 5 μM; FCCP, 5 μM; antimycin A, 10 μM; and rotenone, 2 μM. The OCR was measured using the Seahorse XFe96 extracellular flux analyzer (Andersen et al., 2019; Au - Traba et al., 2016).

### pHluorin live-cell imaging

Transfected MLO-Y4 - pHluorin plasmid cells were used for live-cell imaging. Cells were maintained in a recording medium and images were captured with a Zeiss LSM 810 confocal laser scanning microscope with ZEN imaging software. Transfected cells with good pHluorin signal were first located under a 10X objective and the images were then acquired using a 40X water objective. Images were acquired at a frame size of 512X512 pixels with a time interval of 5 sec between each frame.

### Protein pull-down assay

GST fusion protein containing Cx43 CT and GST control without fusion protein were individually expressed in *E. coli* and purified by binding to glutathione-coupled agarose beads as described on published procedures (Jiang et al., 1994). By using purified GST and recombinant GST-CT fusion protein, we performed the GST pull-down assay (Sambrook and Russell). MLO-Y4 cells transfected with *Atp5j2*- EGFP were cultured in a 10-cm culture plate and cells were collected after 2 days of culturing. Crude cell membrane was prepared and resuspended in a buffer assay medium (20 mM Tris, 50 mM NaCl, pH 7.5) containing nonionic (NP-40, Triton X-100) and zwitterionic (CHAPS). The crude membrane extracts were used in the pull-down experiments. The glutathione agarose beads (Piece, IL) were washed and equilibrated with PBS, aliquoted 50 μL of the beads into two microcentrifuge tubes. The beads were incubated with 20 μg GST-CT or the same amount of GST control for 1 hr at 4°C. An equal amount (1mg) of crude membrane extracts was then added to the tubes containing GST-CT and the GST vials. Incubated the mixture overnight at 4°C. The glutathione agarose beads were washed with PBS five times for 5 min each. An elution buffer (20mM Tris, pH 8.0) containing 10 mM reduced glutathione was used to elute the proteins bound to the beads. Together with the input samples, the eluted fractions were analyzed using 10% SDS–PAGE, and immunoblotting using an anti-GFP antibody (1:2000, #ab290, Abcam, Cambridge MA).

### FRET microscopy

For FRET microscopy, MLO-Y4 cells were transfected with ATP5J2- YFP and Cx43-CFP plasmids. After 48 hrs of culturing, cells were imaged with an LSM- 810 confocal microscope and an argon laser (Carl Zeiss, Thornwood, NY, USA) at excitation wavelength 458 nm for CFP and 514 nm for YFP, and emission wavelength 470- 500 nm for CFP and 530-600 nm for YFP and FRET channels. Confocal images were acquired sequentially in CFP and YFP channels. Identical parameters were used within each channel as well in all test groups. Images were processed using the FRET analysis plugin in NIH ImageJ software.

### Isolation of primary osteocytes from long bone

Primary osteocytes from *Csf-1^+/+^* (WT) and *Csf-1^+/-^* mice were prepared as previously described (Werner *et al*., 2020). Briefly, femurs and tibias were isolated from 21 days male mice with bone marrow removed. Carefully remove the soft tissue attached on the bone surface. Cut the cortical bone into 1 mm pieces and then incubated with 300U/mL collagenase type 1 (Sigma, St. Louis, MO, USA) solution for 3 times, 5 min each, and discarded digests. Then, incubated bone pieces with either 4 mM EDTA or collagenase at 37°C for 30 min twice to remove other cell types. Then incubated the remaining cortical bone with 4 mM EDTA and then with collagenase, collect the fractions after fraction 7. These late digests were filtered through a 100µm cell strainer and were enriched osteocytes. Cultured the primary osteocytes with coated dishes with α-MEM (#12561056, Life Technologies, Carlsbad, CA, USA) supplemented with 5% FBS and 5% BCS (Hyclone, Logan, UT, USA). All animal protocols were performed following the National Institutes of Health guidelines for care and use of laboratory animals and approved by the UT Health San Antonio Institutional Animal Care and Use Committee (IACUC).

### RNA extraction and RT-qPCR

Total RNA was isolated from MLO-Y4 cells using TRIzol reagent (#15596026, Invitrogen, Waltham, MA, USA). cDNA libraby was established by a high-capacity cDNA reverse transcription kit (#4388950, Applied Biosystems, Carlsbad, CA, USA), according to the manufacturer’s instructions. mRNA level was analyzed with two step amplification (94 °C for 10 s, and 60 °C for 30 s, 40 cycles) using a ABI 7900 PCR device (Applied Biosystems, Bedford, MA, USA). The primers for RT-qPCR are: *Sdha*, Forward 5’-GAGATACGCACCTGTTGCCAAG-3’, Reverse 5’-GGTAGACGTGATCTTTCTCAGGG-3’; *Atp5j2*, Forward 5’-CGAGCT GGATAATGATGCGGGA-3’, Reverse 5’-GCAGTAGCTGAAAACCACGTAGG-3’; *Slc25a5*, Forward 5’-ACACGGTTCGCCGTCGTATGAT-3’, Reverse 5’-AAAGCC TTGCTCCCTTCATCGC-3’; *Gapdh*, Forward 5’-CTTCAACAGCAACTCCCAC TCTTC-3’, Reverse 5’-TCTTACTCCTTGGAGGCCATGT-3’.

### Statistical analysis

Statistical analysis was conducted using GraphPad Prism 9 software (GraphPad Software, La Jolla, CA). Two-tailed t test was conducted to the comparison between 2 groups. Multiple group comparisons was done using one-way ANOVA. Two-way ANOVA tests and Tukey multiple comparison tests were used to compare the mean differences between two independent variable groups. The data were presented as mean ± SEM of at least 3 measurements. *p* < 0.05 was designated as a statistically significant difference. \**p* < 0.05; \*\**p* < 0.01; \*\*\**p* < 0.001; \*\*\*\**p* < 0.0001. Images were analyzed using NIH Image J software.

### Materials availability

The newly created plasmids generated in this study are available from the lead contact with a completed materials transfer agreement and without restriction.

### Data availability

All data were included in the manuscript. Source data has been provided.

## Acknowledgment

We thank Professor Yulong Li at Peking University for generously providing the pHluorin plasmid and Dr. Lynda Bonewald at Indiana University for generously providing MLO- Y4 cell line. This work was supported by the National Institutes of Health (NIH) Grants: CA148724 and TL1TR002647 (to F.M.A), and 5RO1 AR072020 (to J.X.J.), and Welch Foundation grant: AQ-1507 (to J.X.J.).

---

**Figure 4-figure supplement 1.**
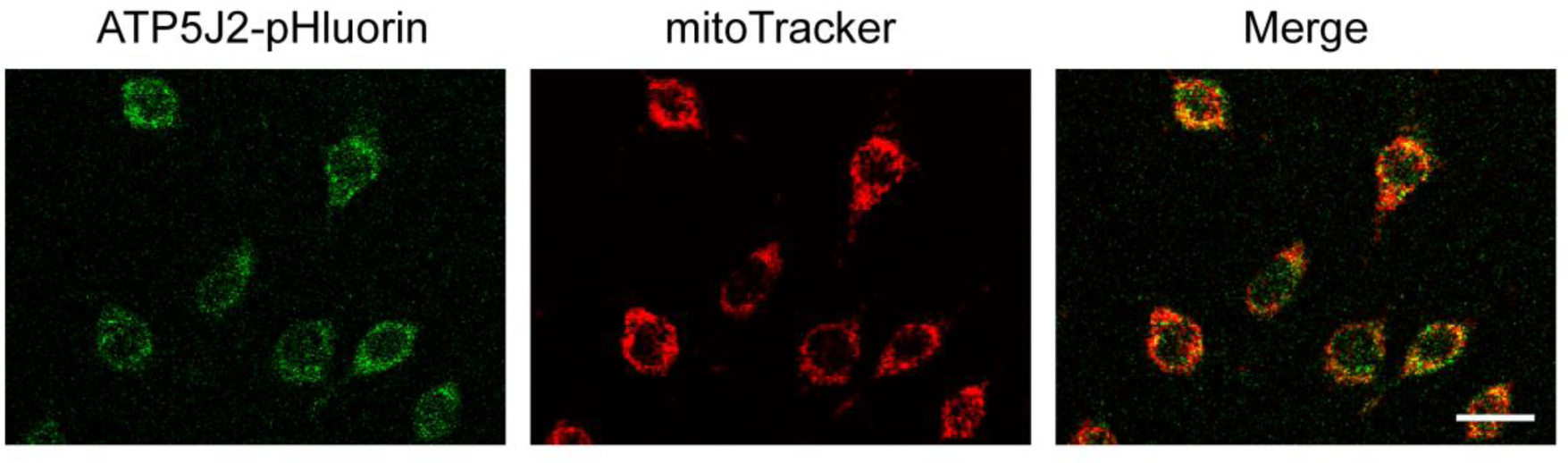
ATP5J2-pHluorin localized on mitochondria. ATP5J2-pHluorin signal was merged with mitoTracker on mitochondria in ATP5J2-mTagBFP- pHluorin transfected MLO-Y4 cells. Scale Bar: 20 µm.

**Figure 5-figure supplement 1.**
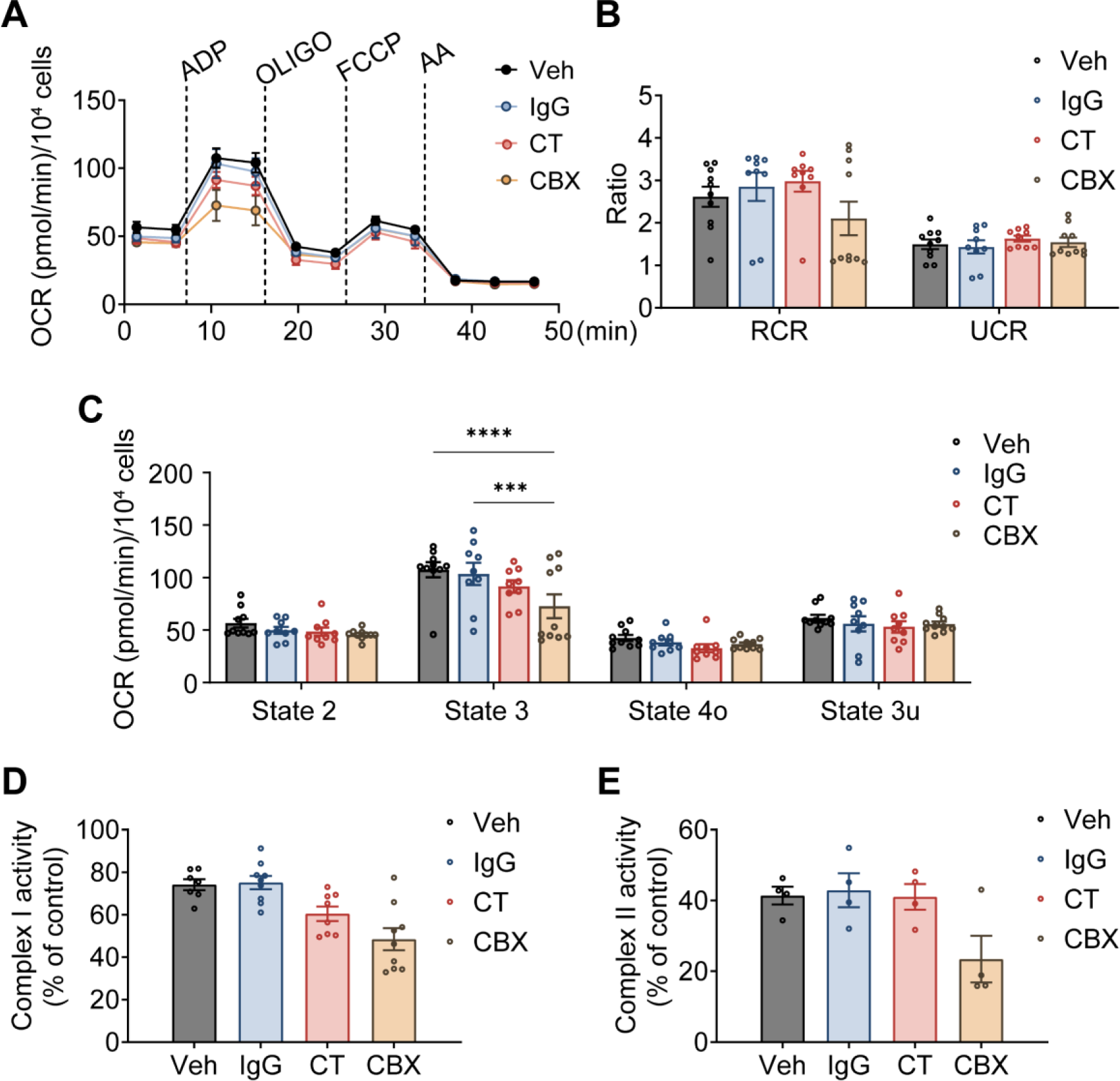
Mitochondrial function was affected under Cx43CT Ab treatment in Cx43 KD MLO-Y4 cells. **(A)** Oxygen consumption rate (OCR) was measured on permeabilized Cx43 KD MLO-Y4 cells, using an XF96e analyser. The sequential injection of mitochondrial inhibitors is indicated by arrows. OLOGO, oligomycin; AA, antimycin A. rabbit IgG, Cx43CT antibody, CBX were pre-added in the assay medium. Basal respiration (state 2), after the injections of ADP (state 3), oligomycin (state 4o), FCCP (state 3u). **(B)** The respiratory control ratio RCR (RCR=state 3/state 4o) and UCR (UCR=State 3u/state 4o), which reflects the mitochondrial respiratory capacity, had no difference between groups. one-way ANOVA test. **(C)** OCR was detected significantly difference in state 3 and state 3u. Values corresponding to the different respiratory states are represented as mean ± SEM (n=15–18 replicates from three independent experiments/group). two-way ANOVA analysis was performed. \**p*<0.05, \*\*\**p*<0.001, \*\*\*\**p*<0.001. **(D-E)** CT antibody and CBX treated group showed the inhibitory effect on complex I and II activity in Cx KD MLO-Y4 cells. Activities of mitochondrial complexes I and II were normalized to the control MLO-Y4 cell line group. \**p*<0.05, \*\**p*<0.01, \*\*\**p*<0.001, \*\*\*\**p*<0.001.

**Figure 6-figure supplement 1.**
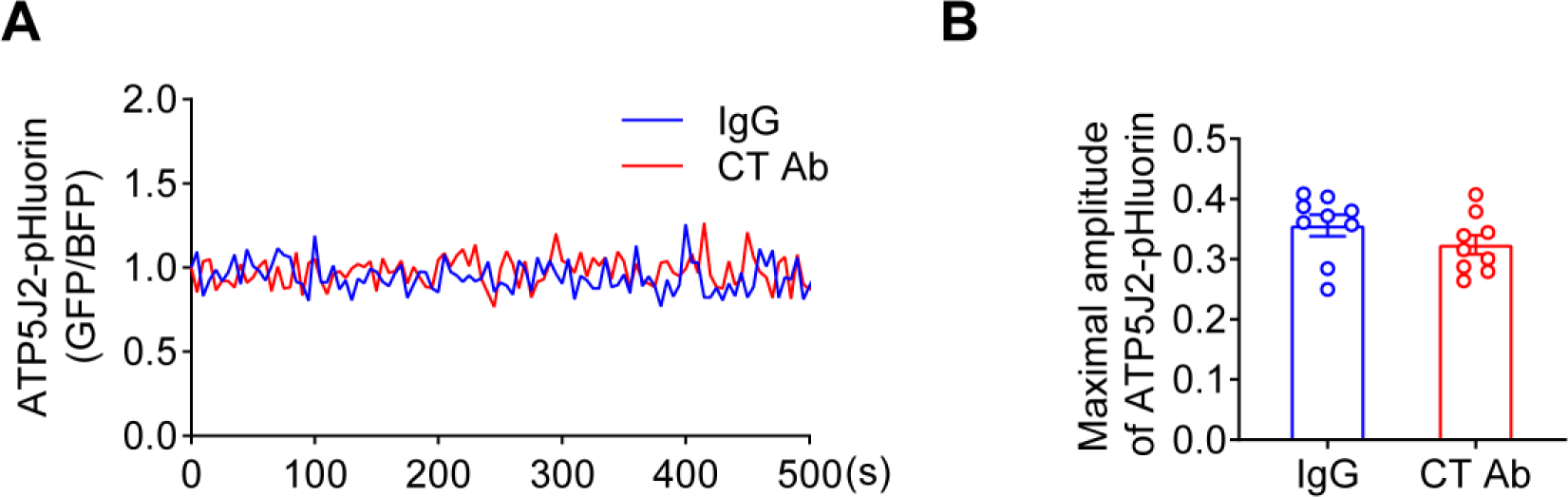
ATP5J2-pHluorin spontaneous oscillation on mitochondria. **(A)** CT antibody showed no effect on ATP5J2-pHluorin signal. Mitochondrial pHluorin signal oscillation in ATP5mf-pHluorin was detected as the same as Cx43-pHluorin in transfected MLO-Y4 cells. **(B)** The maximal amplitude of the oscillation was analyzed with a Two-tailed t-test.

**Figure 7-figure supplement 1.**
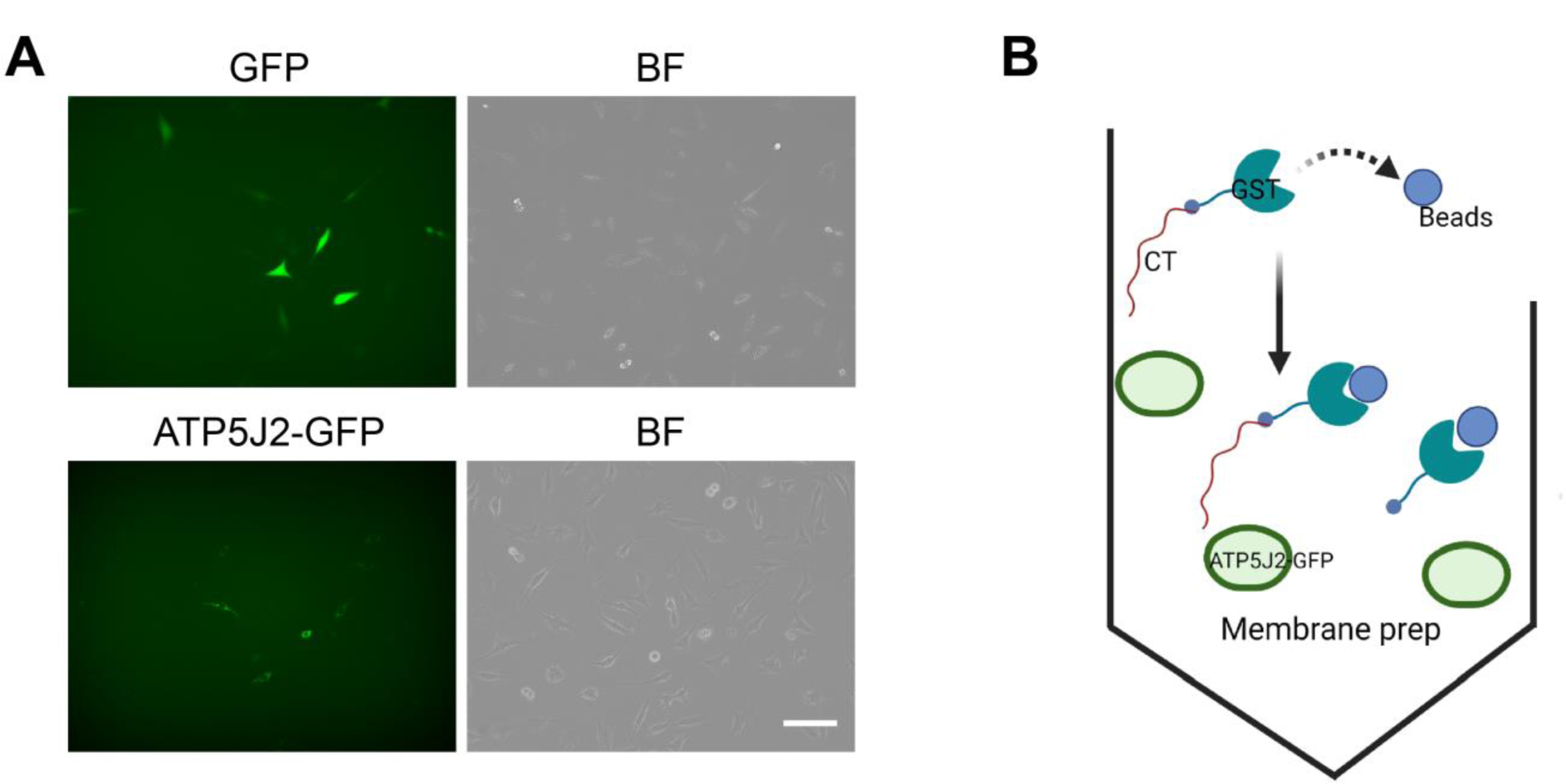
Overexpression of endogenous ATP5J2-GFP in MLO- Y4 cells. **(A)** MLO-Y4 cells were transfected with GFP-N1 or ATP5j2-GFP plasmids. Images were taken in BX-Z100 microscopy with GFP channel and bright field. Scale bar: 100 µm. **(B)** Schematic diagram on GST-pull down assay using transfected MLO-Y4 cells with ATP5J2-GFP and GFP-N1 plasmids as a control.

